# Engineered Adhesion Molecules Drive Synapse Organization

**DOI:** 10.1101/2022.07.18.500465

**Authors:** W. Dylan Hale, T. C. Südhof, R. L. Huganir

## Abstract

In multicellular organisms, cell-adhesion molecules connect cells into tissues and mediate intercellular signaling between these cells. In vertebrate brains, synaptic cell-adhesion molecules (SAMs) guide the formation, specification, and plasticity of synapses. Some SAMs, when overexpressed in cultured neurons or in heterologous cells co-cultured with neurons, drive formation of synaptic specializations onto the overexpressing cells. However, genetic deletion of the same SAMs from neurons often has no effect on synapse numbers, but frequently severely impairs synaptic transmission, suggesting that most SAMs control the function and plasticity of synapses (i.e., organize synapses) instead of driving their initial establishment (i.e., make synapses). Since few SAMs were identified that mediate initial synapse formation, it is difficult to develop methods that enable experimental control of synaptic connections by targeted expression of these SAMs. To overcome this difficulty, we engineered novel SAMs from bacterial proteins with no eukaryotic homologues that drive synapse formation. We named these engineered adhesion proteins ‘Barnoligin’ and ‘Starexin’ because they were assembled from parts of Barnase and Neuroligin-1 or of Barstar and Neurexins, respectively. Barnoligin and Starexin robustly induce the formation of synaptic specializations in a specific and directional manner in cultured neurons. Synapse formation by Barnoligin and Starexin requires both their extracellular Barnase- and Bastar-derived interaction domains and their Neuroligin- and Neurexin-derived intracellular signaling domains. Our findings support a model of synapse formation whereby trans-synaptic interactions by SAMs drive synapse organization via adhesive interactions that activate signaling cascades.

## Introduction

Multicellular organisms rely on cell-adhesion molecules for tissue integrity and intercellular communication. In the vertebrate central nervous system (CNS), distinct classes of cell-adhesion molecules are targeted to synaptic junctions where they organize the composition and performance of the synaptic transmission machinery^1^. Characterization of synaptic adhesion molecules (SAMs) over the past decades revealed an interesting but puzzling pattern: when overexpressed in non-neuronal cells which are then co-cultured with neurons, most SAMs induce the formation of synaptic specializations in the neurons in contact with the non-neuronal cells. When genetically deleted in mice, however, the same SAMs often displayed no apparent function in synapse formation since no deficit in synapse numbers was observed. This paradox is perhaps best exemplified by the well-studied Neuroligin (Nlgn) family of postsynaptic SAMs and the Neurexin (Nrxn) family of presynaptic Nlgn binding partners^2^. Nlgns and Nrxns are single-pass transmembrane proteins that were identified as a trans-synaptic adhesion complex linking cell-adhesion to intracellular scaffolding molecules on either side of the synaptic cleft^3,4^. Overexpression of Nlgns or Nrxns in non-neuronal cells potently induce in co-cultured neurons the formation of pre- or post-synaptic specializations onto these cells, raising the possibility that Nlgns and Nrxns might stimulate synaptic assembly^5–8^. Moreover, overexpression of Nlgns in cultured neurons enhanced synaptic contacts onto overexpressing cells^9–11^.

These results led to a view of synapse formation that relied on Nlgn and Nrxn interactions to drive the assembly of the synaptic machinery at points of close membrane apposition. This model was bolstered by multiple lines of evidence demonstrating that other classes of synaptic molecules, such as those that mediate synaptic vesicle exocytosis, were largely dispensable for synapse formation, suggesting that synapse formation occurs in the absence of neurotransmitter signaling^12–15^. However, increasingly sophisticated genetic approaches have revealed that deletion of Nlgn family members, either alone or in combination, yield at best a minor reduction in synapse numbers^16–19^. The same pattern is observed with Nrxns^20–22^ and is now known to be the case for an increasing number of SAMs^23–32^. We estimate that at least 20 SAMs distributed over 10 gene families follow this pattern of driving synapse organization when overexpressed in non-neuronal cells that are co-cultured with neurons, but that are not essential for synapse formation in an in vivo setting. In contrast to these groups of SAMs, two families of adhesion-GPCRs, BAIs and latrophilins, and their ligands were found to be essential for synapse formation in vivo^33–38^. Viewed together, these results suggest that SAMs fall into two classes, those that ‘make’ a synapse such as Latrophilins and Bai’s, and those that ‘shape’ a synapse, such as Nlgns and Nrxns^1^.

However, even SAMs that induce synapse formation in contacting neurons do not act autonomously but require a precise matching complement of pre- and postsynaptic interacting SAMs^36^.This requirement could account for the specificity of synapse formation but renders facile manipulations of synapse formation difficult. To develop new genetic tools that enable artificial generation of synapses between neurons that normally do not engage in synaptic interactions, we engineered SAMs containing tightly interacting bacterial proteins fused to the C-terminal sequences of Nrxn3β and Nlgn1 as targeting and signal-transduction components. We show that in cultured neurons, these artificial synaptogenic SAMs potently induce formation of synapses, validating the overall concept.

## Results

### Nlgn1 and Nlgn2 but not Nlgn3 induce presynaptic organization

We began the development of genetic tools for manipulating synapse number by first investigating the properties of Nlgn proteins that, when overexpressed, facilitate synapse formation. We started by assessing the three main Nlgn family members (Nlgn1, Nlgn2, and Nlgn3) for differential ability to induce synapse formation when overexpressed in cultured hippocampal neurons. Previous attempts to discern differences in Nlgn-induced presynaptic organization have been complicated by the possibility that Nlgn family members assemble into homo- or hetero-dimers and that the Nlgn dimer is likely the functional unit for Nlgn-Nrxn interaction (**Fig 1*A***)^39–41^. Therefore, overexpression of a single Nlgn family member may drive the other Nlgn family members expressed endogenously to the cell surface and complicate the interpretation of structure-function analyses.

**Figure 1.**
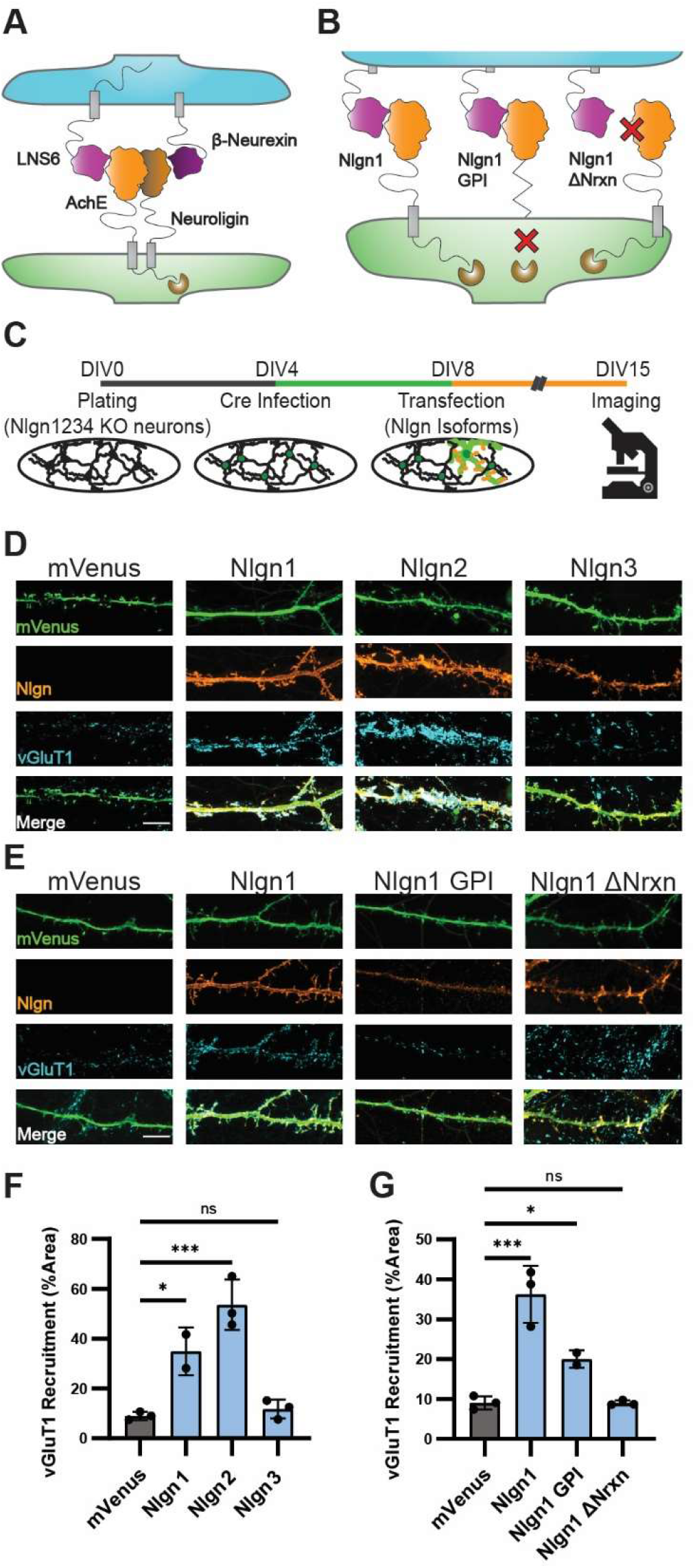
High-affinity neurexin binding is required for Nlgn-induced synapse organization. **A)** Cartoon illustrating Nlgn-Nrxn adhesion complex showing the postsynaptic (green) Nlgn AchE domain (orange) in complex with the presynaptic (blue) Nrxn1β LNS6 domain (purple). **B)** Cartoon of Nlgn1 mutants Nlgn1 GPI lacking the TM and intracellular domain and Nlgn1ΔNrxn failing to interact with Nrxn1β. **C)** Experimental timeline showing hippocampal cultures produced at DIV0, infected with Cre lentivirus at DIV4, and transfected with Nlgn cDNAs at DIV8. Cells were fixed for imaging at DIV15. **D)** Nlgn1 and Nlgn2 overexpression but not Nlgn3 (orange) induce accumulation of vGluT1 (blue) onto transfected dendrites (green). **E)** Nlgn1 GPI weakly induces accumulation of vGluT1 while Nlgn1-ΔNrxn does not meaningfully induce vGluT1 accumulation. **F)** Quantification of (D). One-way ANOVA with Dunnet’s multiple comparison correction (n=2 for Nlgn1, otherwise n=3). **G)** Quantification of (E). One-way ANOVA with Dunnet’s multiple comparison correction (n=2 for Nlgn1 GPI otherwise n=3). (*=p<0.05; ***=p<0.001) Scale bars = 10µm.

To circumvent this issue, we utilized a transgenic mouse line with the genes encoding Nlgn1,-2,-3 flanked by loxP sites (floxed)^42^ and the gene encoding the final Nlgn, Nlgn4 constitutively deleted^43^ (Jamain et al., 2008).In the absence of Cre-recombinase, neurons prepared from these animals lack Nlgn4, but upon Cre delivery the remaining three Nlgn genes are deleted, resulting in Nlgn1,-2,-3,-4 quadruple knockout cultures (NlgnqKO^16,44^).

To assess the relative ability for Nlgn1,-2, and -3 to induce presynaptic organization, we first generated cultures from postnatal day 0 (p0) NlgnqKO mice, and infected these cultures at DIV4 with lentivirus for expression of Cre recombinase expression under the control of a synapsin promoter to produce cultures with a Nlgn-null background (**Fig 1*C***). On DIV10, cultures were transfected with mVenus cDNA either alone or in combination with Nlgn1,-2, or -3 cDNA. At DIV 15, cultures were fixed and stained for the presynaptic marker vGluT1 to mark excitatory presynaptic specializations (**Fig 1 *D***). As expected, Nlgn1 overexpression increased the density of presynaptic specializations onto overexpressing dendrites as compared to mVenus transfection alone. Nlgn2 overexpression also significantly increased the density of presynaptic specializations (**Fig 1*D, F***), in agreement with some prior reports (Linhoff et al., 2009) and contrary to others^11^. Consistent with our previous observations^19^, we found that Nlgn3 overexpression in Nlgn-null cultures did not increase the density of presynaptic specializations compared to mVenus expression alone (**Fig 1*D, F***). We therefore concluded that Nlgn family members exhibit a differential ability to induce synapse formation when overexpressed.

### Neurexin binding is required for Nlgn-induced presynaptic organization

Given the high homology between Nlgn family members, the differential presynaptic induction of Nlgn3 compared to the other two family members was striking. Despite the high degree of conservation among Nlgns at the primary amino acid level, some differences have been observed, including differential synaptic localizations^45–48^, differential affinities for binding to presynaptic neurexins, Nlgn3 binding to Nrxn1β being nearly an order of magnitude weaker than that of Nlgn1 or Nlgn2^49^, and differential functions in that deletion of Nlgn1 only impaired excitatory but not inhibitory synaptic transmission, whereas deletion of Nlgn2 only suppressed inhibitory but not excitatory transmission^19^. Thus, we hypothesized that the difference in affinity may underlie the differential presynaptic induction observed between Nlgn1/-2 and Nlgn3.

To imitate this difference in affinity, we generated a ‘Nlgn1ΔNrxn’ construct that contains the LNDQE mutation which is known to disrupt the binding interface between Nlgn1 and Nrxn1β^39^. As before, we overexpressed this construct at DIV10 into Cre-expressing hippocampal neurons from NlgnqKO animals and monitored vGluT1 levels on the dendrites of the overexpressing neurons (**Fig1*B, C***). Presynaptic organization onto Nlgn1ΔNrxn-expressing dendrites was indistinguishable from that of cells expressing mVenus alone, indicating that neurexin binding is critical for the ability of Nlgn1 to induce presynaptic organization (**Fig. 1*E, G***). To screen for other extracellular features that might explain the ability of Nlgn1 to induce presynapse formation, we tested all eight Nlgn1 splice variants by overexpression in cultured neurons (*Fig. S1*). No statistically significant differences emerged between splice variants.

This finding suggests that Nlgn1 overexpression in neurons does not simply drive synapse organization by boosting the intracellular interaction of Nlgn1 with scaffolding molecules, such as PSD95^50^, thereby driving localization of synaptic scaffolding components and by extension other SAMs to the site of dendritic contact, which could then proceed to induce presynaptic differentiation. Our findings contradict this model and raise the question whether the transmembrane region and intracellular sequences of Nlgn1 are required for driving the organization of synapses. To address this question, we generated a glycosylphosphatidylinositol- (GPI-) tethered Nlgn1 extracellular domain by fusing the first 693 amino acids of Nlgn1 to a 2X HA-tag followed by the NCAM GPI-anchoring sequence^8^.This construct results in a Nlgn1 acetylcholinesterase-homology domain anchored to the external leaflet of the plasma membrane with no extracellular stalk domain or intracellular domain (**Fig. 1*B*)**. While overexpressing this construct in NlgnqKO neurons resulted in expression much weaker than that observed with our other Nlgn1 constructs, it still induced presynaptic specializations to accumulate on overexpressing dendrites, demonstrating that the extracellular domain of Nlgn1 is sufficient to induce presynaptic organization (**Fig. 1*E, G***). Together these data provide clear evidence that the induction of presynaptic organization by Nlgn1 overexpression only requires the extracellular domain interaction of Nlgn1 with presynaptic neurexins.

### Engineered adhesion molecules derived from a bacterial RNAse complex

Our data indicate that adhesion between Nlgn1 and presynaptic neurexins is required for induced presynaptic organization, raising the question whether any extracellular adhesion interaction targeted to synapses might facilitate synapse organization. While Nlgn1 is the best characterized postsynaptic binding partner for presynaptic neurexins, neurexins bind to an array of other molecules, both in *cis* and in *trans*^2,51,52^. Moreover, Nlgns bind in *cis* to postsynaptic MDGAs which may influence their interaction with neurexins^53–55^, and likely bind to additional SAMs because the Nrxn-binding-deficient mutant of Nlgn1 still exhibits specific synaptic functions^56^. These extensive interaction networks make comprehensive mutagenesis screens prohibitively complex and time intensive. We therefore sought instead to eliminate *cis* or *trans* interactions by engineering our own adhesion domains to replace the adhesive globular domains of Nlgns and Nrxns, thereby eliminating most extracellular interactions in a single stroke.

We began by searching for protein domains with no known homologs in the mammalian CNS and with a high interaction affinity. We settled on a well-studied protein pair from the bacterium *Bacillus amyloliquefaciens*, the small RNAse barnase and its inhibitor barstar^57^. In addition to possessing one of the highest affinities of interaction of known protein pairs, barnase and barstar have no homologs in any vertebrate system. To convert these proteins into adhesion molecules, the coding sequence for the LNS6 domain of Nrxn3β was replaced with a gene fragment encoding the inhibitor barstar to create a new molecule that we dubbed ‘Starexin’ (**Fig. 2*A*, Fig. *S2***). Nrxn3β was chosen for the construction of Starexin to be able to compare it to previous neurexin structure-function analyses using Nrxn3β^8^. Similarly, we removed the acetylcholinesterase homology domain (AchE, **Fig. 2*A*, Fig. *S2***) coding sequence from our Nlgn1 cDNA and replaced it with a gene fragment encoding barnase to create a molecule that we dubbed ‘Barnoligin’. While Starexin expressed well in cells, expression of Barnoligin alone in either HEK cells or neurons produced 100% cell death in <24 h, reflecting the known toxicity of the barnase molecule in the absence of the barstar inhibitor. We hypothesized that this toxicity was due to the RNAse activity of barnase. To address this, we screened several previously characterized mutations in barnase thought to reduce RNAse activity while preserving interaction with barstar^58^. Mutating Barnoligin to carry the H102Q mutation previously identified to reduce the RNAse activity resulted in a protein that was expressed well with no observable effects on cell health. We therefore proceeded with H102Q Barnoligin, hereafter referred to simply as ‘Barnoligin.’

**Figure 2.**
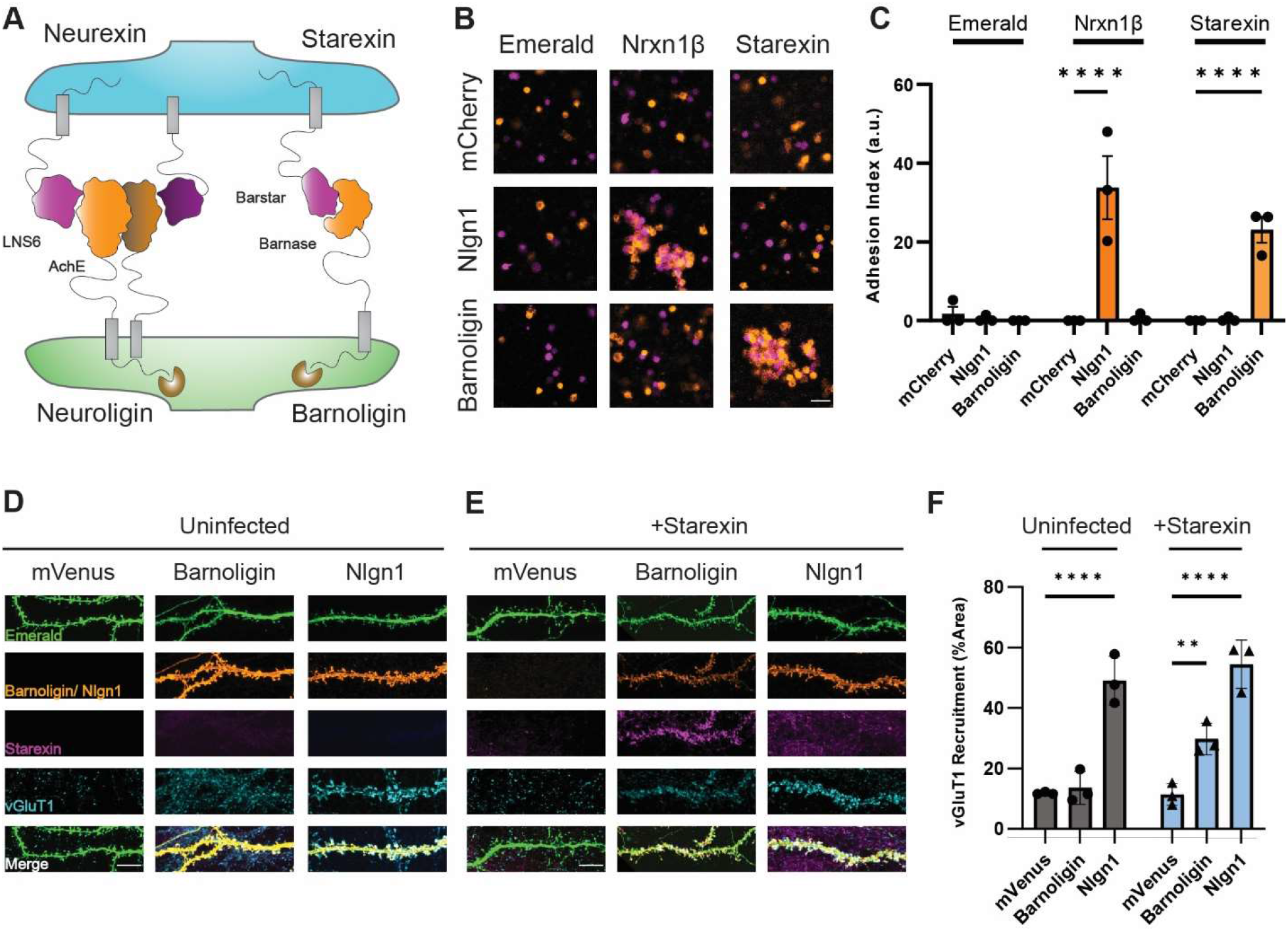
Barnoligin and Starexin – engineered adhesion molecules derived from a bacterial RNAse complex. **A)** Cartoon depicting the design of presynaptic Starexin (right side, purple) and postsynaptic Barnoligin (right side, orange). Cartoon shows the barnase and barstar protein domains replacing the Nlgn1 AchE (left side, orange) and Nrxn3β LNS6 (left side, purple) domains, respectively. Note that in contrast to Nlgn1, Barnoligin is known not to form homodimers. **B)** HEK293F cells grown in suspension and expressing mCherry (pseudo-colored orange) or Emerald (pseudo-colored purple) do not aggregate (top left panel), but when cells expressing Nlgn1 (orange) and cells expressing Nrxn1β (purple) are mixed, aggregates form as a result of cell-adhesion (center panel). When cells expressing Barnoligin (orange) and Starexin (purple) are mixed, they form adhesive aggregates like Nlgn1 and Nrxn1β. Scale Bar = 50µm. **C)** Quantification of (B) 2-way ANOVA with Dunnet’s multiple comparisons (n=3). **D)** Barnoligin overexpression in uninfected neurons fails to induce vGluT1 accumulation. **E)** In contrast, Barnoligin overexpression(orange) in Starexin-expressing neurons drives Starexin (purple) accumulation along transfected dendrites (green) which in turn generates vGluT1 accumulation (blue). Scale bar = 10µm. **F)** Quantification of (D) and (E). Two-way ANOVA with Dunnet’s multiple comparisons (n=3). (**\*\***=<0.01; **\*\*\*\***=<0.0001)

To test whether these molecules could function as an adhesion pair akin to Nlgns and Nrxns, we transfected either Barnoligin, Starexin, Nlgn1, or Nrxn1β into suspension-cultured HEK-293F cells alongside fluorescent mCherry or Emerald reporter constructs. After 48 h, we mixed cells expressing different adhesion molecules together and agitated the cells for 1 hour to determine if our engineered proteins drove the formation of an adhesion complex. As expected, cells expressing Nlgn1 + mCherry adhered with cells expressing Nrxn1β + Emerald forming large clumps that were absent when cells expressing Nlgn1 + mCherry or Nrxn1β + Emerald were mixed with cells expressing Emerald or mCherry alone (**Fig. 2*B***). Cells expressing Barnoligin + mCherry or Starexin + Emerald adhered together and formed large clumps similar to Nlgn1 and Nrxn1β aggregates (**Fig. 2*B***). However, cells expressing Barnoligin + mCherry did not adhere to cells expressing Nrxn1β + Emerald nor did cells expressing Nlgn1 + mCherry adhere to cells expressing Starexin, indicating that our engineered adhesion proteins are specific for their engineered partners (**Fig. 2*C***).

### Barnoligin-Starexin adhesion induces presynaptic organization

After observing Barnoligin and Starexin form a specific adhesion complex in mammalian cells, we tested whether these molecules could induce synapse organization in cultured neurons. To this end, we prepared hippocampal cultures from newborn WT mice and transfected Barnoligin at DIV10 under the control of the human synapsin promoter into either uninfected neurons or neurons infected at DIV4 with lentiviruses expressing V5-tagged Starexin under the control of the human synapsin promoter. Overexpression of Barnoligin via transfection in uninfected neurons did not noticeably change the organization of presynaptic terminals compared to overexpression of mVenus alone (**Fig. 2*D***). However, overexpression of Barnoligin via transfection in neurons infected with Starexin lentiviruses yielded three striking effects. First, accumulation of Starexin was observed on Barnoligin-transfected dendrites, indicating the reconstitution of the adhesion complex in cultured neurons (**Fig. 2*E***). Secondly, the excitatory presynaptic marker vGluT1 similarly accumulated onto Barnoligin-transfected dendrites at a level significantly above that of cells expressing mVenus alone (**Fig. 2*E*, 2*F***). Finally, transfection of Nlgn1 induced presynaptic organization irrespective of whether neurons expressed virally-induced Starexin, indicating that the expression of Starexin alone does not diminish the synapse organizing potential of other SAMS in culture (**Fig. 2*F***).

### Barnoligin-Starexin adhesion directs compartment-specific synaptic organization

The fact that Barnoligin and Starexin can induce synapse organization despite their adhesion domains being completely foreign to the mammalian nervous system raises interesting possibilities about how synaptic organization proceeds. Given that Barnoligin and Starexin lack the endogenous globular extracellular domains native to the parent molecules Nlgn1 and Nrxn3β, it seems improbable that synapse organization requires additional *cis* or *trans* interactions from these domains. These data suggest that the adhesion formed by the Nlgn1 AchE domain and the Nrxn3β LNS6 domain is functionally sufficient to drive synapse organization and does not require other extracellular molecules in the complex for synaptic organization to proceed. In addition to adhesion, are other features of SAMs required for synaptic organization? Or is synapse organization the programmed response of neurons to adhesion?

To begin to address these issues, we asked if the synaptic organization induced by Barnoligin and Starexin was directional or compartment specific. That is, we tested whether the remaining Nlgn1 and Nrxn3β sequences present in Barnoligin and Starexin were sufficient to specify their organizational function to pre- or postsynaptic compartments, respectively. To answer this question, we relied on a heterologous co-culture assay in which non-neuronal cells expressing SAMs are cultured with mature cultured neurons^6,9,59^. This approach is cleaner than transfection in cultured neurons because heterologous cells do not express common synaptic markers such as vGluT1 or Homer1. Therefore, any synaptic organization observed is solely a result of the cultured neurons responding to the SAMs on the heterologous cell surfaces. We infected cultured WT mouse neurons at DIV4 with our lentivirus driving V5-tagged Starexin expression under the control of a synapsin promoter. At DIV 15, HEK293 cells expressing the fluorescent cell fill Emerald either alone or along with Barnoligin or Nlgn1 were co-cultured with uninfected neuron cultures or cultures infected with Starexin lentivirus. After 48 hours of co-culture with uninfected neurons, presynaptic organization by HEK cells expressing Barnoligin was indistinguishable from those expressing Emerald alone (**Fig. 3*A***). However, when co-cultured with neurons expressing Starexin, Barnoligin-expressing HEK cells potently induced vGluT1 accumulation similar to that induced by Nlgn1 expression (**Fig. 3*B***). HEK cells expressing Nlgn1 potently induced the organization of vGluT1 irrespective of whether the cells were infected with Starexin lentivirus (**Fig. 3*A-B*, 3*E***).

**Figure 3.**
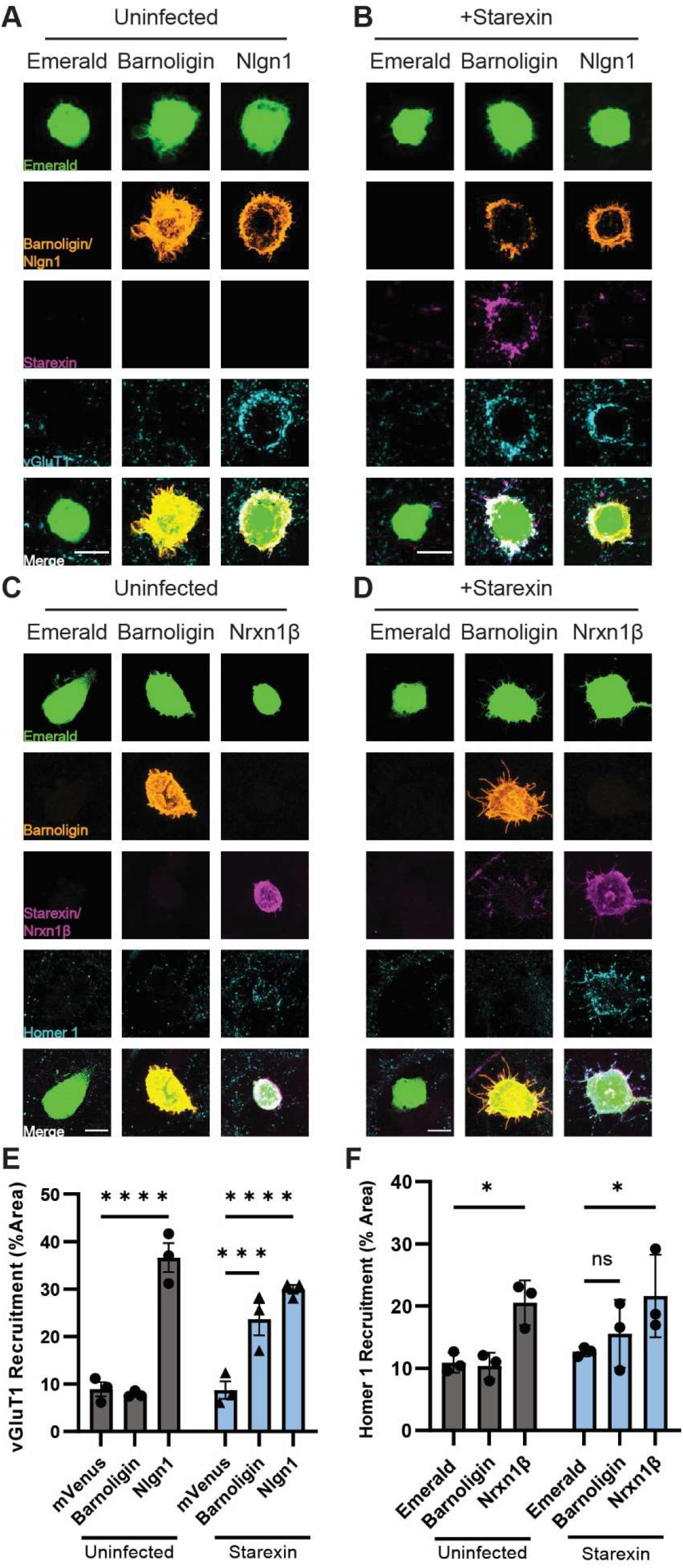
Starexin specifically facilitates the organization of presynaptic, but not of postsynaptic, specializations onto Barnoligin-expressing HEK293 cells. **A)** Barnoligin-expressing HEK cells co-cultured with uninfected neurons show no vGluT1 accumulation on their cell surface as compared to cells expressing Emerald alone. **B)** When co-cultured with neurons expressing Starexin, cells expressing Barnoligin accumulate vGluT1 on the cell surface similar to cells expressing Nlgn1. **C)** Barnoligin-expressing HEK cells do not accumulate Homer-1 positive postsynaptic specializations when co-cultured with uninfected neurons, **D)** and this is also true when neurons are expressing Starexin. Note that although Barnoligin on the surface of HEK cells accumulates Starexin-positive processes, no increase in Homer1 is observed. **E)** Quantification of A (gray) and B (blue). **F)** Quantification of C (gray) and D (blue). Statistical comparisons made with 2way ANOVA with Dunnet’s multiple comparison correction. (*=p<0.05; ***=p<0.001; ****=p<0.0001).

Similarly, HEK cells expressing the presynaptic molecule Nrxn1β induced a significant increase in post-synaptic organization in both infected and uninfected cultured neurons as seen by staining for the excitatory postsynaptic marker Homer1 (**Fig. 3*C*-*D***). However, in contrast to presynaptic organization, HEK cells expressing Barnoligin failed to have any appreciable effect on Homer1 organization in neurons expressing Starexin (**Fig 3*D*, 3*F***).

To determine if the observed compartment-specificity was true in the opposite orientation, we generated lentivirus for the expression of HA-tagged Barnoligin under the control of a synapsin promoter. After 48 hours of co-culture HEK cells expressing V5-Starexin showed no effect on post-synaptic organization in uninfected neurons (**Fig. 4*A***). However, in neurons infected at DIV4 with Barnoligin lentivirus, co-culture of HEK cells expressing Starexin resulted in dramatic induction of Homer1 organization after 48 hours (**Fig. 4*B*, 4*E***). Nrxn1β induced postsynaptic organization irrespective of whether the neurons expressed Barnoligin (**Fig. 4*B*, 4*E***). In contrast, HEK cells expressing Starexin failed to induce presynaptic organization as seen by vGluT1 staining irrespective of whether co-cultured neurons were expressing Barnoligin (**Fig. 4*C*-*D*, 4*F***). Together, these data indicate that Barnoligin and Starexin induce synaptic organization in a directional, compartment-specific manner.

**Figure 4.**
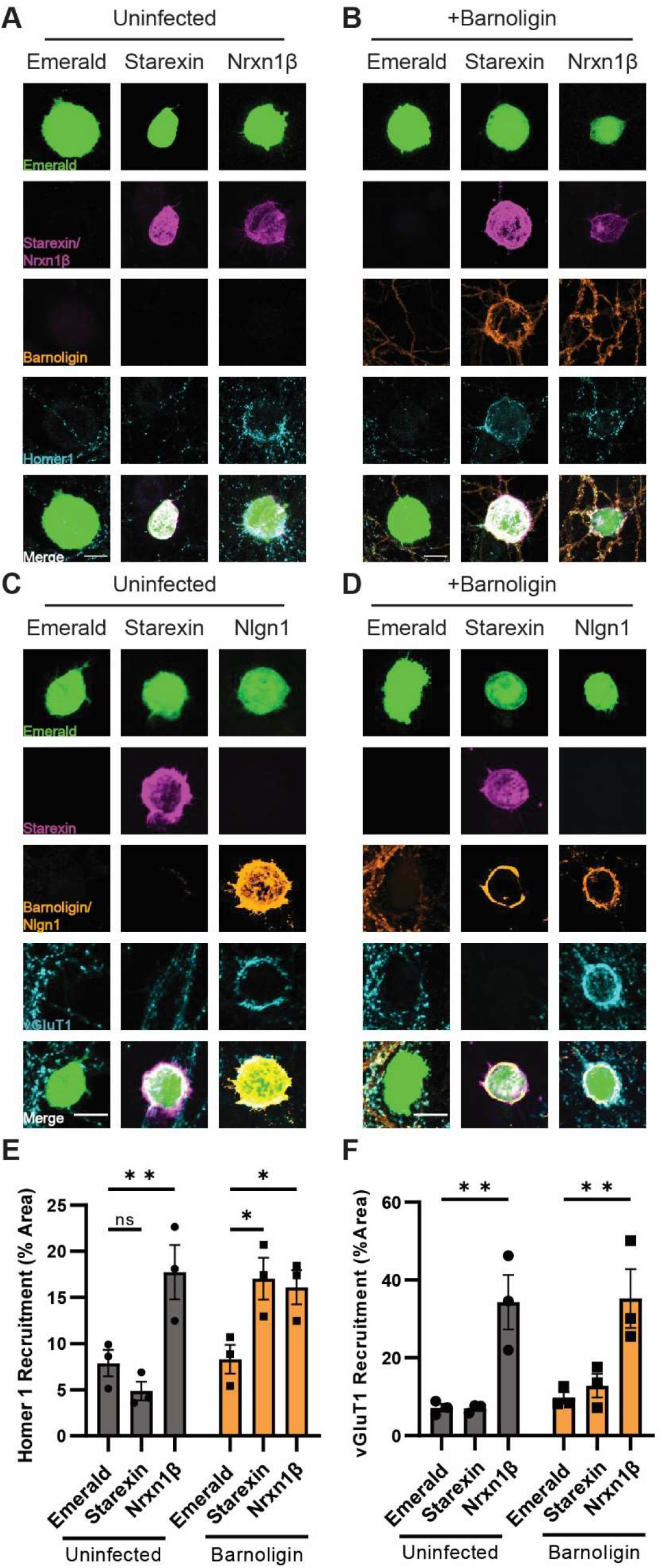
Barnoligin specifically facilitates the organization of postsynaptic, but not of presynaptic, specializations onto Starexin-expressing HEK293 cells. **A)** Starexin-expressing HEK293 cells co-cultured with uninfected neurons fail to induce the accumulation of Homer-1 positive postsynaptic specializations. **B)** In neurons expressing Barnoligin, Starexin HEK cells induce striking Barnoligin accumulation onto the cell surface accompanied by an accumulation of Homer1-positive postsynaptic specializations. Infection with Barnoligin does not affect the ability of Nrxn1β to induce postsynaptic organization. **C)** Starexin-expressing HEK 293 cells co-cultured with uninfected neurons also fail to accumulate vGluT1-positive presynaptic specializations onto the cell surface. **D)** Starexin-expressing HEK cells also fail to induce vGluT1 accumulation when co-cultured with neurons expressing Barnoligin. Barnoligin expression does not inhibit Nlgn1 from inducing presynaptic organization. **E)** Quantification of A (gray) and B (orange). **F)** Quantification of C (gray) and D (orange). Statistical comparisons made with 2way ANOVA with Dunnet’s multiple comparison correction. (*=p<0.05; **=p<0.01).

### Starexin-induced presynaptic organization requires the Nrxn3β intracellular domain

While the directionality of Barnoligin and Starexin adhesion-induced synaptic organization suggests that each molecule engages in specific signaling to facilitate synaptic organization, there is still a possibility that synaptic organization is a generic neuronal response to adhesion and that the directionality observed with Barnoligin and Starexin is an accidental property, and not produced by differences in intracellular signaling. To address this possibility, we generated a GPI-anchored version of Starexin, dubbed Starexin-GPI. Attaching the Starexin extracellular domain to the external leaflet of the cell membrane via a GPI-anchor allowed us to eliminate any intracellular signaling that the intracellular domain of Nrxn3β contributed to Starexin (**Fig. 5*A***). As previously tested with Starexin, Starexin-GPI was capable of specifically forming an adhesion complex with Barnoligin as measured by HEK-293F aggregation (**Fig. 5*B*-*C*;** for full comparison, see **Fig S3**) We therefore generated lentiviruses for the expression of Starexin-GPI under the control of a synapsin promoter as with full-length Starexin. To test whether Starexin-GPI could induce presynaptic organization similar to full-length Starexin, we plated HEK293T cells expressing Barnoligin onto DIV15 cultured WT hippocampal neurons that were either uninfected, infected with Starexin lentivirus, or infected with Starexin-GPI lentivirus. As before, Barnoligin-expressing HEK cells failed to induce vGluT1 organization in uninfected neurons (**Fig. 5*D***) but vigorously induced vGluT1 organization in Starexin-expressing neurons (**Fig. 5*E***). While Barnoligin-expressing HEK cells demonstrably formed a strong adhesion complex with Starexin-GPI expressing neurons, no vGluT1 accumulation was immediately obvious (**Fig. 5*F***). However, careful quantification revealed that significantly more vGluT1 was present on and around Barnoligin HEK cells co-cultured with neurons expressing Starexin-GPI as compared with uninfected neurons (**Fig. 5*G***). This effect was significantly less potent than that seen when Barnoligin HEK cells were co-cultured with neurons expressing full-length Starexin (**Fig. 5*G***) and lacked the characteristic ‘halo’ of vGluT1 around Barnoligin HEK cells observed in the Starexin expressing neurons (**Fig. 5*H***). We quantified the number of HEK cells that showed these vGluT1 ‘halos’ from a random sample of Barnoligin-expressing HEK-293 cells co-cultured with neurons expressing either Starexin or Starexin-GPI and found that while the overwhelming majority of Barnoligin-expressing HEK cells co-cultured with Starexin-expressing neurons showed a vGluT1 ‘halo’, this characteristic effect was demonstrably absent from nearly all Barnolgin-expressing HEK cells co-cultured with neurons expressing Starexin-GPI (**Fig. 5*I***). We therefore conclude that while the Nrxn3β intracellular domain of Starexin is required for synapse organization, Starexin-GPI incidentally increases the amount of vGluT1 in contact with Barnoligin-expressing cells owing to increased axonal contact as a product of the strong adhesion produced by Barnoligin and Starexin.

**Figure 5.**
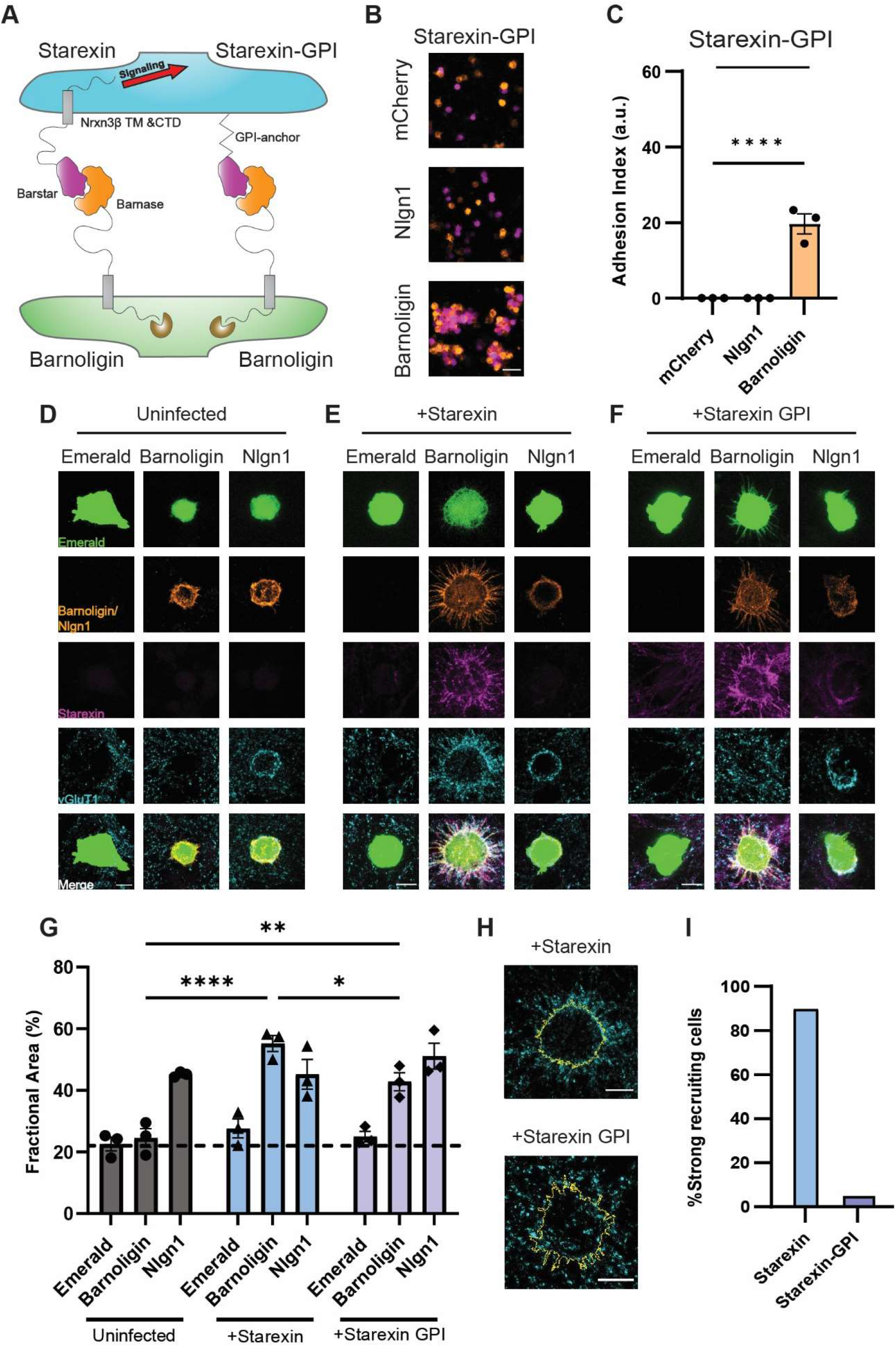
Intracellular signaling is required for Starexin to direct presynaptic organization. **A)** Cartoon depicting the design of the Starexin-GPI construct. **B)** Suspension HEK293F cells expressing Starexin GPI (purple, all panels) specifically form an adhesion complex only with cells that are expressing Barnoligin (orange, bottom panel) and not mCherry alone or Nlgn1 (both orange, top and middle panels respectively). **C)** Quantification of B. **D)** As before, Barnoligin-expressing HEK cells do not induce presynaptic accumulation in uninfected neurons. **E)** Starexin expressed in neurons directs synapse organization to the surface of Barnoligin-expressing co-cultured HEK cells. **F)** Neuronally-expressed Starexin GPI accumulates on the surface of Barnoligin-expressing HEK cells as Starexin does, but the characteristic accumulation of vGluT1 is absent. **G)** Quantification of D, E, F (Gray, blue, lavender, respectively). Although characteristic ‘halos’ of vGluT1 are absent from Starexin GPI-expressing neurons, careful quantification reveals a significant increase in presynaptic specializations co-incident with Barnoligin-expressing HEK cells. **H)** Close-up detail of two Barnoligin-expressing HEK cells co-cultured either with Starexin (top) or Starexin-GPI (bottom). Cell outlines are shown in yellow. **I)** Quantified fraction of Barnoligin-expressing cells co-cultured with Starexin or Starexin GPI-expressing neurons demonstrating characteristic recruitment ‘halos’. Statistical comparisons in C, G made with 2way ANOVA with Dunnet’s multiple comparison correction. (*=p<0.05; ***=p<0.001; ****=p<0.0001).

**Figure 6.**
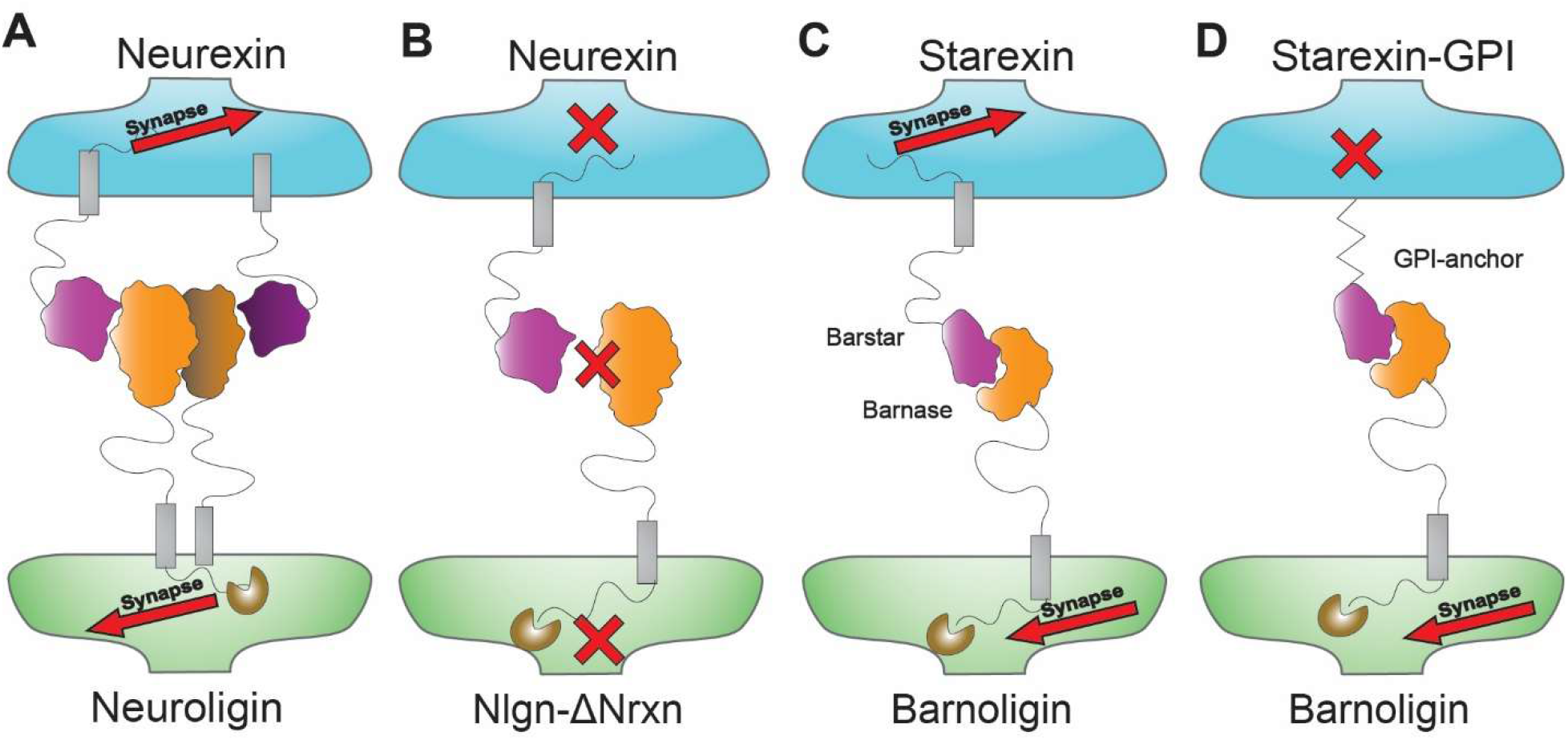
Summary of results – adhesion and intracellular signaling are both required for synapse organization. A) Adhesion by Neurexins and Neuroligins can drive synapse organization which is blocked B) when the binding interface between Nlgn1 and Nrxns is disrupted. C) Barnoligin and Starexin can drive synapse organization through adhesion but synapse organization is blocked D) by the replacement of the Nrxn3β intracellular domain with a GPI anchor.

## Discussion

### Adhesion as a fundamental property of synapse organization

Understanding the mechanisms that drive synapse organization is one of the most fundamental challenges in cellular neuroscience. For decades evidence has accumulated that SAMs play a fundamental role in the organization and alignment of synaptic machinery, but specifically testing the role of adhesion as a physical force contributing to synapse formation has been challenging. While mouse genetics has facilitated the inactivation of entire SAM gene families^17,20,21,25,42^, the ability to eliminate *all* synaptic adhesion would require a complete accounting of every SAM in the vertebrate CNS and the technological means to disable them all simultaneously. Absent this, our approach of generating novel adhesion molecules elegantly demonstrates that adhesion between complimentary neuronal membranes likely represents at least one requirement for the organization of synaptic machinery.

This may help to explain the extraordinary diversity among SAMs capable of inducing synaptic organization through overexpression in cultured neurons^1,2^. Extracellular domains of SAMs need only share the common feature of forming a functional adhesion complex of sufficient strength in *trans* to induce synapse organization. Therefore, diversity in extracellular domains allows for the specificity required for neurons to precisely wire to their proper synaptic partners. However, this only partially resolves the puzzling relationship of SAMs with synapse organization as our data reveal that our newly engineered adhesion molecules also require intracellular sequences derived from SAMs to be fully effective at organizing synapses (**Fig. 5**). Therefore adhesion, while seemingly essential, is not sufficient for the induction of synaptic organization but also requires intracellular signaling once the adhesion complex is formed. In the case of adhesion GPCRs like the Bai’s and latrophilins, this downstream signaling likely involves both G protein-dependent and G protein-independent signaling cascades^60^. In the case of Nrxns and Nlgns, downstream signaling is less clear but may converge on canonical protein kinase pathways, including the PKA and DLK→JNK signaling pathways. One explanation of how SAM overexpression might drive synapse organization is that runaway signaling by kinases downstream of overexpressed SAM adhesion could aberrantly activate targets that are normally reserved for activation by GPCRs, thereby imitating the synaptogenic signaling of adhesion GPCRs. Another possibility is that SAMs drive localization of adhesion GPCRs to sites of aberrant adhesive contact through interactions with intracellular scaffolding molecules, thereby signaling the formation of synapses through conventional adhesion GPCR activation - although the mechanism of GPCR activation in this model is unclear. In either case, our approach of replacing the adhesive domains of SAMs with barstar and barnase is an effective and generalizable strategy for probing the relative contributions of adhesion to synaptogenic signaling.

### Barnoligin and Starexin reveal principles of cell-adhesion

Our successful engineering of Barnoligin and Starexin is fortuitous. Before we engineered an adhesion complex with these two molecules, it was unclear what differentiated an adhesion molecule from a generic transmembrane receptor. Our intuition and experience working with SAMs led us to believe that adhesion molecules required a few key properties to function: 1) Adhesion molecules must be transmembrane or otherwise membrane-anchored; 2) Adhesion molecules must have complimentary extracellular domains that allowed for the formation of homomeric or heteromeric complexes in *trans*; and 3) The strength of adhesion is roughly analogous to the KD of biochemical interaction and that this KD must surpass some as yet undefined threshold for adhesion. The barnase and barstar proteins met the third criterion before we began, and we designed Barnoligin and Starexin in such a way that they would meet the remaining two criteria. That this approach succeeded suggests that our intuition about adhesion molecules was at least partially correct and that future attempts can help refine our understanding of what properties govern cell-adhesion. The success of Barnoligin and Starexin as adhesion molecules and their validation of our adhesion principles may also explain the evolution of SAMs from disparate evolutionary lineages. The adhesion domain of Nlgns is evolutionarily derived from the acetylcholinesterase family of enzymes^61,62^. Similarly, the adhesion domain of Barnoligin was engineered from the enzyme barnase – a molecule with no known adhesion function. By co-opting an intracellular RNAse and its inhibitor as adhesion domains of our own design, we may have mirrored an evolutionary pattern in which proteins that were refined through evolution for one purpose may have later been repurposed as adhesion molecules. We suspect that the principal limitation for whether an extracellular protein can be co-opted as an adhesion molecule is whether a high-affinity binding partner can be generated, by evolution or design, to couple to its extracellular surface. These insights could prove useful in both the discovery of unconventional adhesion molecules as well as in future designs for engineered adhesion molecules.

### Barnoligin and Starexin as tools for manipulating synaptic inputs

Barnoligin and Starexin were conceived and designed by incorporating insights and questions from our experience studying synapse organization by SAMs. Beyond insights into cellular neuroscience Barnoligin and Starexin represent unique tools for continuing to probe synaptic connectivity. The most interesting question is whether Barnoligin and Starexin could be used to alter connectivity in an intact brain by viral induction of Starexin in projection cells with simultaneous delivery of Barnoligin to postsynaptic cells downstream of Starexin-expressing axons. Such an experiment would alter synaptic connectivity in a way that has not been previously achieved, representing an important opportunity to understand how alterations in circuit connectivity may affect the behavior of entire organisms. This approach will be particularly powerful when coupled with cell-type specific expression, allowing for the reweighting of inputs in neuronal circuits. Such a targeted and specific manipulation of circuits will be of tremendous benefit to the study of diverse neurological and neuropsychiatric diseases. Beyond neuroscience, Barnoligin and Starexin could be used as a platform in many tissue systems to probe outstanding questions about how adhesion-based signaling or perturbed adhesion influences cellular behavior. Collectively, our findings are both technology and insight – channeling decades of work on the fascinating question of how SAMs affect synapse organization into a tool that can broadly address questions of how adhesion affects cellular behavior and tissue organization.

## Materials and Methods

### Animal Husbandry

All mouse lines used in this study were maintained in accordance with institutional guidelines and protocols for humane animal treatment. Complete details regarding the NlgnqKO mouse line were published previously (Zhang et al., 2018).

### DNA Constructs and Vectors

Nlgn constructs were cloned from mouse cDNA into the pDisplay vector in frame with the N-terminal IgK signal sequence and HA tag but out of frame of the PDGFR tail. Nlgn1 splice variants were produced via inverse PCR of A1,A2,B ‘all in’ Nlgn1 to delete each splice site either alone or in combination. Nrxn1β was cloned similarly but the HA tag was replaced with a V5 via PCR. Cloned cDNAs of Nlgn1 and Nrxn1β were identical to those found at: Nlgn1 (GenBank: NM_138666.4), Nrxn1β (GenBank: NM_001346960.2). Nlgn1-GPI was produced by amplification of the Nlgn1 cDNA corresponding to amino acids 46-690 with homologous primer tails homologous to the NCAM GPI-anchoring plasmid reported previously^8^. Barnase and barstar from *Bacillus amyloliquifasciencs* cDNAs were codon-optimized and synthesized with homologous ends to existing Nlgn1 or Nrxn3β plasmids and ligated with In-fusion enzyme (Takara Bio). A graphical summary of the design of major fusion constructs used can be found in ***Fig S2***. All plasmids were fully commercially sequenced prior to experimental application. Sequences for the Barnoligin and Starexin plasmids used in this study were deposited with Genbank: pCMV5 Barnoligin (ON997589), pCMV5 Starexin (ON997590), pCMV5 Starexin-GPI (ON997591), pFSW Barnoligin (ON997592), pFSW Starexin (ON997593), pFSW Starexin-GPI (ON997594).

### Neuronal culture

P0 hippocampal mouse neurons were generated as previously described^19^. Neurons were plated at 1.5×10^5^ cells/mL into 24-well plate wells in Serum Media (MEM (Life Technologies), 5.4% Fetal Bovine Serum (Atlanta Biologicals), 2mM L-Glutamine (Life Technologies,), 2.5% B-27 (Gibco), with D-Glucose (Sigma) added up to 6.6mM) onto Matrigel-coated number 0 coverslips (Carolina Biologicals). After 24 hrs, 80% of media was replaced with Growth Media (Neurobasal (Life Technologies) with 2% B-27 supplement and 2mM L-Glutamine). 50% of culture media was changed every 4 days beginning DIV4. Cells were harvested at DIV15 for neuron overexpression experiments and DIV 17 for co-culture.

### HEK 293T Cell Culture

Human Embryonic Kidney (HEK) 293T cells were obtained from (ATCC) and expanded for 2 passages before use in experiments. Cells were maintained in Dubelcco’s Modified Eagle Medium (DMEM (Life Technologies)) with 10% FBS.

### Lentivirus Production

Cre and ΔCre Synapsin Lentiviruses were obtained by providing pFSW-Cre and pFSW-ΔCre plasmids to the Stanford University Neuroscience Gene Vector and Virus Core who then produced Syn-Cre and Syn-ΔCre viruses. Starexin-V5, starexin-GPI-V5, and barnoligin-HA viruses were produced externally from pFSW-Starexin, pFSW-Starexin-GPI, and pFSW-Barnoligin-HA plasmids (Janelia, Ashburn, VA). Each virus was empirically tested for infectious titer before being employed in experiments.

### Calcium Phosphate Transfection

For HEK cell transfection, 1μg of each plasmid of interest was mixed with 10.664μL of 2M CaCl2 and the total volume was brought up to 86μL with sterilized MilliQ H2O. The resulting precipitate was incubated for 10 minutes before being added dropwise to cells in a 6-well plate well. Cells were incubated for at least 24 hours before being employed in co-culture experiments.

### Cell Aggregation Assay

Freestyle 293-F cells (Thermo Fisher) were grown in 40mL of Freestyle Expression medium (Thermo Fisher) in 125mL culture flasks shaking at 125rpm at 37C and 5% CO2 to a density of 2×10^6^ cells/mL with a viability of 90% or greater. Cells were then transfected with polyethylineamine (PEI, 40,000MW, Polysciences) using 24μg of PEI and 8μg total plasmid per condition. 48 hours post-transfection, cells from different flasks were mixed 1:1 in uncoated 12-well plates and shaken in a culture incubator at 125rpm for 1hr. Live confocal microscopy was used to assess aggregation. The fraction of total fluorescence observed in aggregates larger than a single cell size was reported as the adhesion index.

### Immunostaining

Coverslips with cultured neurons or HEK 293T cells were fixed with 4% Paraformaldehyde in 1X PBS for 30 minutes at 4C. Coverslips were washed three times with 1mL of 1X PBS before being permeabilized with 0.2% Triton X-100 in 1X PBS for 5 minutes at room temperature and blocked with 5% BSA solution in 1X PBS for 30 minutes to 1 hour. Primary immunostaining was done overnight (12-16 hours) at 4C with the appropriate antibody dilution in 5% BSA solution. After primary immunostaining, cells were washed three times with 1X PBS before being incubated with secondary antibody solution in 5% BSA solution for 1 hour at room temperature. Coverslips were then washed three times with 1X PBS solution before being washed once with deionized H2O and mounted onto charged glass slides into a drop of Fluoromount G (Southern Biotech). Slides were allowed to dry overnight before imaging. Antibodies and dilutions used in this study are as follows: Chicken anti-HA (Aves Labs, 1:1000); Mouse anti-HA (Covance (now Biolegend) 1:1000); Mouse anti-V5 (Invitrogen, 1:500); Guinea Pig anti-vGluT1 (Synaptic Systems 1:1000); Mouse anti-GAD65 [GAD6] (Abcam 1:1000); Rabbit anti-Homer1b/c [135 304] (Synaptic Systems, 1:1000); Cy3 Donkey anti-chicken (Jackson Immuno, 1:1000); Goat anti-Guinea Pig Alexa Fluor 647 (Invitrogen, 1:1000); Goat anti-mouse IgG2a Dylight 405 (Jackson immune, 1:1000); Goat anti-mouse Alexa Fluor 567 (Invitrogen 1:1000).

### Confocal Microscopy

Slides were imaged at 63xwith a Nikon A1 confocal using NIS elements or Zeiss LSM880 using the Zen software. Acquisition settings were kept constant for every sample and condition within each experimental replicate. Samples were imaged blind to treatment, using mVenus or Emerald signal to identify regions of interest. Optical slices were collected at 0.2μM to give optimal resolution in the Z dimension.

### Image analysis and statistics

Images were analyzed in FIJI ((NIH, USA; RRID: SCR_003070) in a semi-automated fashion using macros by masking transfected neuronal dendrites or HEK cell bodies and measuring the fractional area of each mask covered by the synaptic stain of choice. Data analysis was conducted blind to experimental treatment. Experimental treatments were unblinded when fully processed quantitative data was entered into GraphPad Prism for statistical analysis and plotting.

## Notes

### Competing Interest Statement

The authors have declared no competing interest.

