## Supplement for "Engineered Adhesion Molecules Drive Synapse Organization"

**SUPPLEMENTAL DATA**

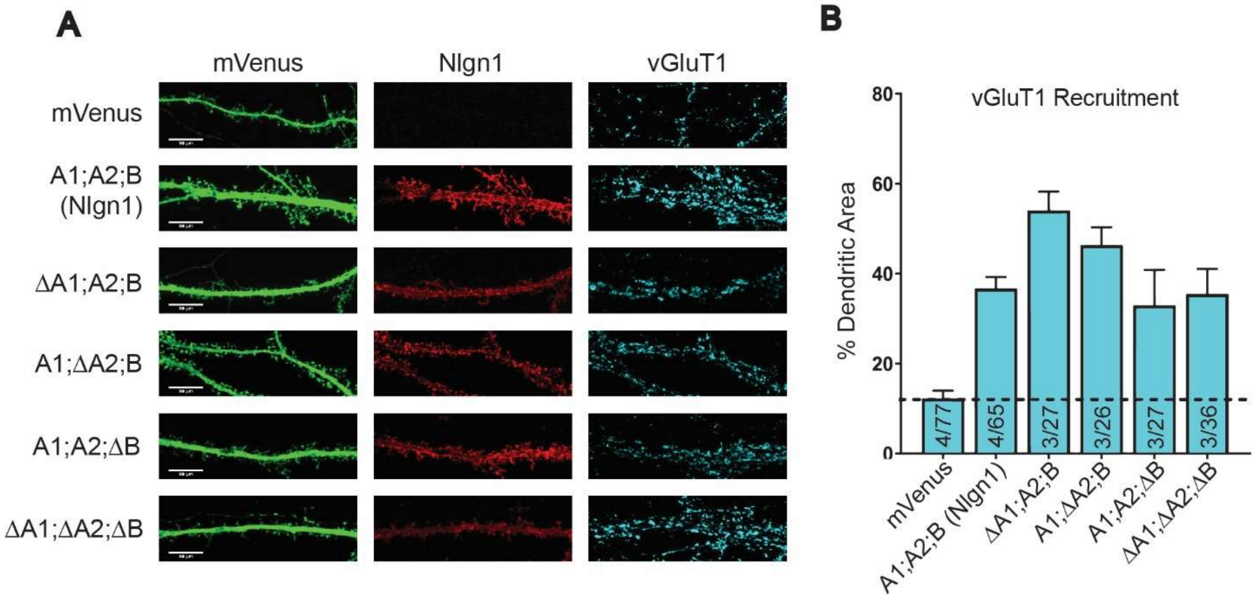


**Figure S1. Neuroligin1 Splicing does not impair synapse-organization. A)** Representative images showing vGluT1 staining on dendrites overexpressing Nlgn1 constructs with splice sites 'in' or splice sites removed (Δ) either individually or in combination. **B)** Quantification of the excitatory presynaptic marker vGluT1 accumulation on dendrites overexpressing Nlgn1 splice forms. Nlgn1 is alternatively spliced at two sites, A and B. The A site has two different sequences that can be spliced in either independently or together A1 and A2. All eight splice variants were tested but high variability and the large number of statistical comparisons made quantitative comparisons difficult. Qualitative evaluation of overexpressed Nlgn1 splice forms showed no obvious differences in the ability to drive excitatory or inhibitory presynaptic marker accumulation.

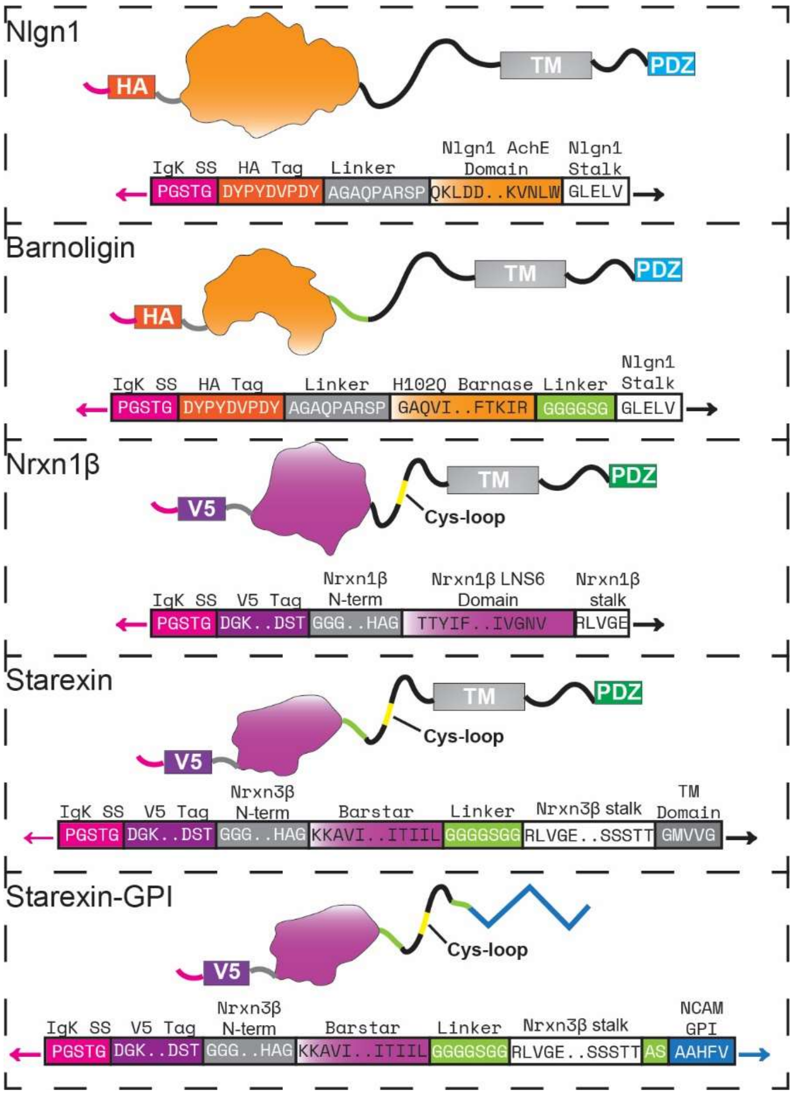


**Figure S2. Design of Barnoligin, Starexin and Starexin GPI relative to Nlgn1 and Nrxn1β proteins.** Top of each panel – cartoon representation of each protein showing globular domains and unstructured regions. Bottom of each panel – amino acid sequences at critical junctions. Elipses indicate that more than three amino acids are interspersed between the specific residues notated. To ensure consistent expression of each construct, the endogenous signal peptides of Nlgn1 and Nrxn1β were replaced with the Immunoglobulin-kappa signal peptide (Igk SS). Affinity tags are located on the N-terminus immediately following the signal sequence and then a short linker sequence before beginning the globular domains. The position of the Nrxn1β cysteine loop as well as the positions of the Nrxn3β cysteine loops present in starexin constructs are indicated in yellow. Pink arrows indicate that the polypeptide continues to the N-terminus, black arrows indicate that the polypeptide continues to the c-terminal PDZ domains and the blue arrow indicates that the polypeptide continues to the c-terminus of the *M. musculus* NCAM 120 GPI-anchoring motif. Sequences used to construct: Nlgn1 (GenBank: NM_138666.4), Nrxn1β (GenBank: NM_001346960.2). Full sequences submitted to GenBank corresponding to: pCMV5 Barnoligin (ON997589), pCMV5 Starexin (ON997590), pCMV5 Starexin-GPI (ON997591), pFSW Barnoligin (ON997592), pFSW Starexin (ON997593), pFSW Starexin-GPI (ON997594).

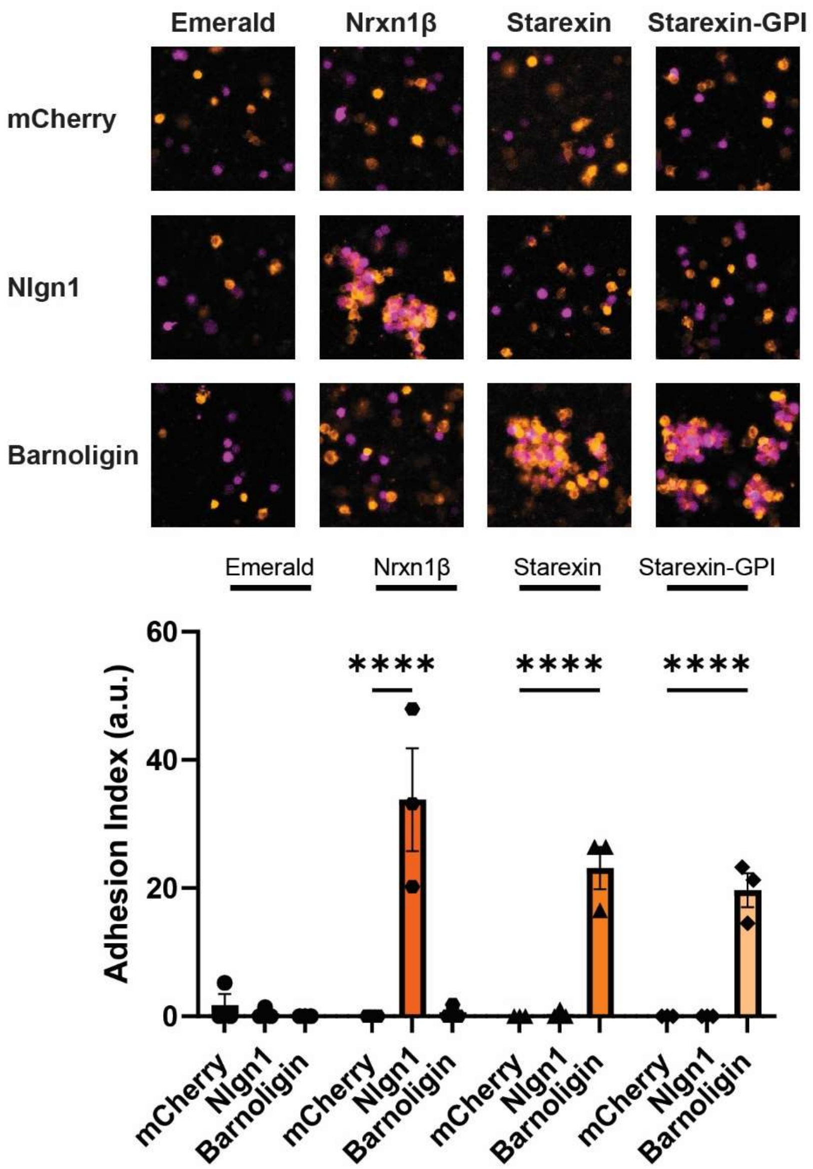


**Figure S3. Full comparison of adhesion molecules in the aggregation assay.** Cell aggregates were observed when Nlgn1 and Nrxn1β were combined and when Barnolign and either Starexin or Starexin GPI were combined. Compared to Starexin, Starexin-GPI produced similar levels of aggregation with Barnoligin. Statistical comparisons were made with 2-way ANOVA with Dunnet’s Multiple comparisons. ****$p<0.0001$.

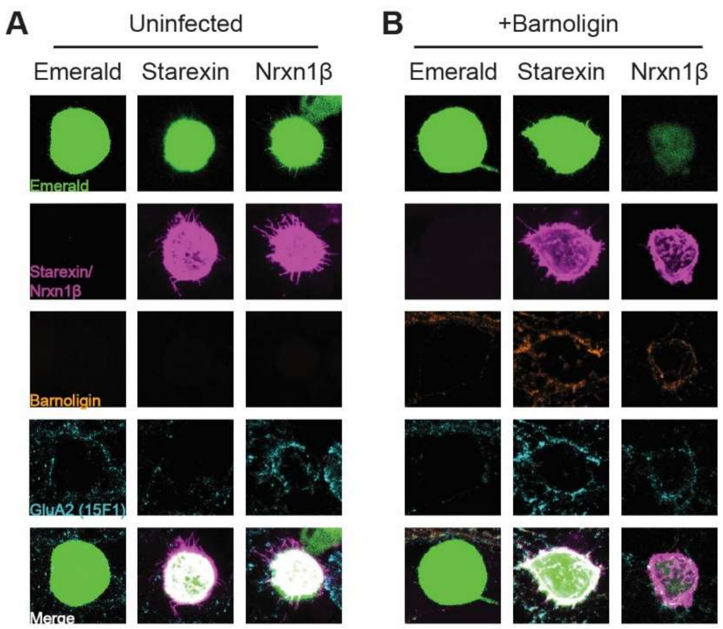


**Figure S4. Barnoligin-Starexin organizes postsynaptic receptors. A)** Co-culture of Starexin-expressing HEK 293 cells with cultured hippocampal neurons did not result in accumulation of AMPAR subunit GluA2 revealed by surface staining. **B)** However, co-culture of Starexin-expressing HEK293 cells with hippocampal neurons expressing Barnoligin did result in a pattern of GluA2 accumulation around the HEK cell. Nrxn1β drove accumulation of GluA2 irrespective of whether Barnoligin was expressed in neurons.
